# Interactive design and validation of antibody panels using single-cell RNA-seq atlases

**DOI:** 10.1101/2024.10.15.618479

**Authors:** Matthew Watson, Simon Latour, Golnaz Abazari, Michael J Geuenich, Eunice Poon, Ruonan Cao, Miralem Mrkonjic, Alison P. McGuigan, Hartland W. Jackson, Kieran R. Campbell

**Affiliations:** Lunenfeld-Tanenbaum Research Institute, Sinai Health System, Toronto ON M5G 1×5, Canada; Department of Molecular Genetics, University of Toronto, Toronto ON M5T 3A1, Canada; Data Sciences Institute, University of Toronto, Toronto ON M5G 1Z5, Canada; Institute of Biomedical Engineering, University of Toronto, Toronto ON M5S 3G9; Department of Laboratory Medicine & Pathobiology, University of Toronto, Toronto ON M5S 1A8, Canada; Ontario Institute of Cancer Research, Toronto, ON M5G 1M1, Canada; Department of Chemical Engineering and Applied Chemistry, University of Toronto, Toronto, ON, M5S 3E5; Vector Institute, Toronto, ON M5G 1M1, Canada; Department of Statistical Sciences, University of Toronto, Toronto ON M5S 3G3, Canada; Department of Computer Science, University of Toronto, Toronto ON M5S 1A8, Canada

## Abstract

Single-cell RNA-sequencing holds promise for identifying novel markers of cellular variation for antibody-based technologies. However, antibody panel design is often difficult due to multiple experimental and biological constraints. We introduce Cytomarker, an interactive platform enabling human-in-the-loop design of antibody panels from single-cell transcriptomic data. We use Cytomarker to spatially profile human mammary tissue subpopulations and develop a novel antibody screening approach to validate granular subpopulation predictions across >3.5M cells.

## Main

Multiple technologies enable efficient multiplexed quantification of protein expression using antibody panels for profiling or enrichment with spatial and single-cell resolution^1–4^. These have driven discoveries of both novel cell states or functions and more recently cellular location or organization predictive of patient outcomes^5,6^. However, most of these technologies only quantify up to tens of proteins simultaneously, necessitating careful design of the antibody panel that best captures unique cell subsets or broad biological variation in the tissue context of interest. Therefore, most existing work uses well-established markers of cell lineage (e.g. immune cell subsets) state (e.g. hypoxic) or function (e.g. cytotoxic) to build out such panels.

Concurrently, international initiatives such as the Human Cell Atlas^7^ (HCA) and Human Tumor Analysis Network^8^ have, or plan to, generate whole transcriptome single-cell expression profiles for virtually all cell types across both normal and diseased tissues. This raises the exciting possibility of data-driven panel design that best captures unique cell subtypes or encompasses the cellular composition of a tissue by searching transcriptome-wide for targets. While several methods have been proposed to find an optimal marker set given a single-cell RNA-seq (scRNA-seq) dataset^9–12^, these one-pass methods have multiple drawbacks for practical panel design. Not all markers suggested will have validated antibodies, and not all antibodies will be validated for the specific application of interest. Similarly, experimentalists likely already possess multiple antibodies they would prioritize if “good enough” at capturing cellular heterogeneity as a cost-saving mechanism. Finally, human supervision of panel design enables the ability to prioritize cell populations of interest while maintaining canonical markers that align with current best-practice or may not be represented well at the transcriptome level^13^.

Given these issues, we sought to create an interactive human-in-the-loop platform that best exploits both computational and human contributions to antibody panel design using scRNA-seq. Our solution — Cytomarker (**Fig. 1A** and www.cytomarker.ai) — can take either a custom annotated scRNA-seq uploaded by the user in a variety of common formats^14–16^ or access scRNA-seq datasets from a multitude of tissues and cell types from the HCA. Cytomarker next suggests an initial panel of existing antibody reagents that best captures cellular heterogeneity, either using existing methods^10^ or using a custom cell-type aware algorithm (**Methods**). Cytomarker then produces several interactive visualizations to enable the user to assess the suitability of the panel, including heatmaps and violin plots of marker genes, as well as UMAP^17^ visualizations computed using both the full scRNA-seq dataset and the currently selected panel. In addition, Cytomarker introduces a machine learning procedure to quantitatively score a given panel, enabling the user to rationally compare and improve panels.

**Figure 1.**
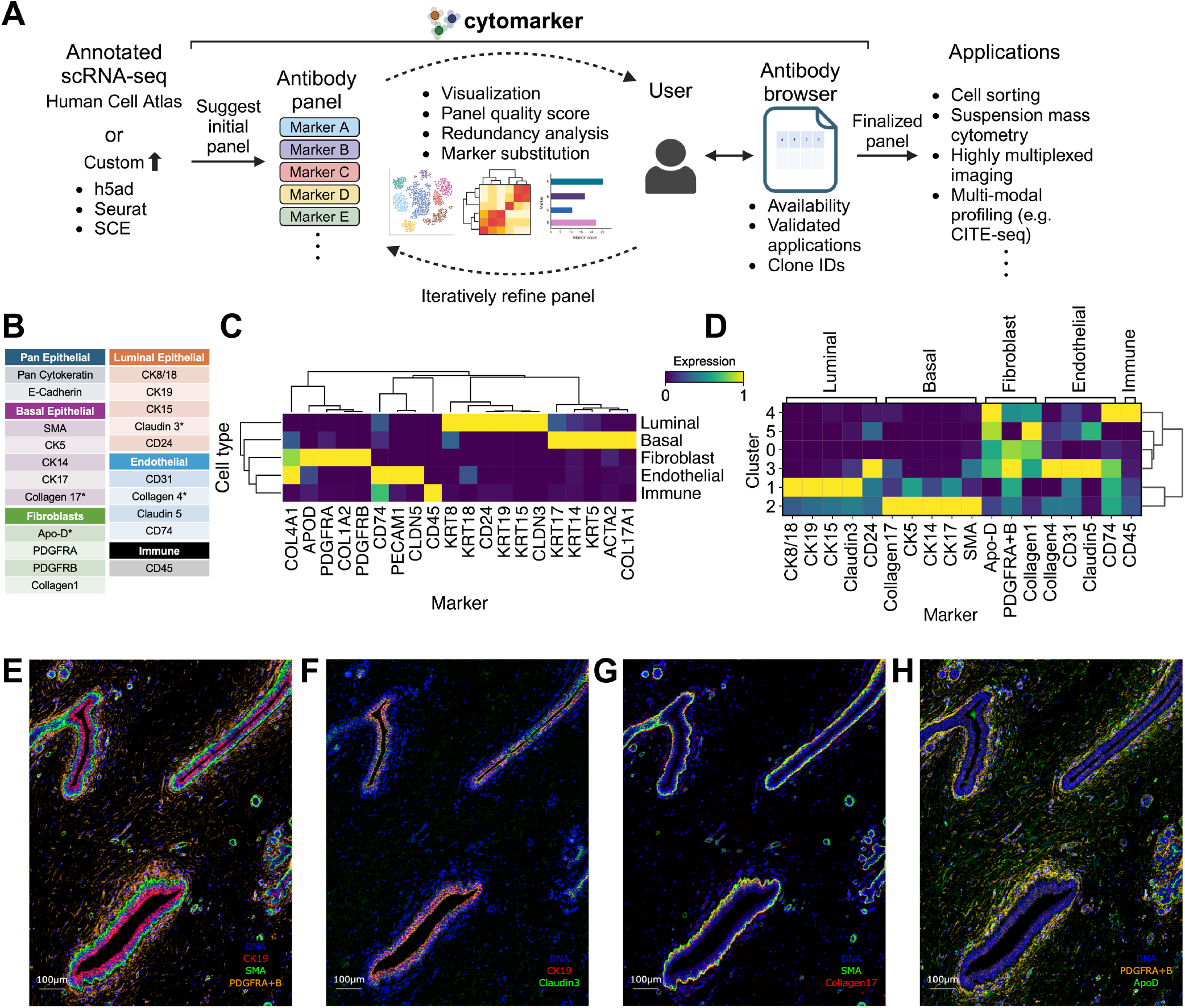
Cytomarker iteratively learns antibody panels from scRNA-seq atlases. **A** The cytomarker workflow takes custom or atlas scRNA-seq data, suggests an initial panel preserving biological heterogeneity, before enabling an iterative process of panel refinement. Finally, targets are linked to a database of antibodies covering a number of applications. **B** The antibody panel for profiling healthy mammary tissue designed using Cytomarker with scRNA-seq. * denotes putatively novel targets. **C** Cluster specific expression profiles in the scRNA-seq. **D** Cluster-specific expression profiles from subsequent profiling with IMC. **E** Representative tissue region showing canonical markers for luminal epithelial (CK19), basal epithelial (SMA), and fibroblast (PDGFRA-B) populations. **F-H** Novel subpopulation markers Claudin3, Collagen17, ApoD on the same region of tissue.

Next, Cytomarker enables multiple routes for a user to iteratively improve a given panel. Users may add Cytomarker-suggested markers to improve the ability of a panel to capture a given cell type or state. Additionally, if a user wishes to reduce the panel size either for cost savings or to add other markers targeting orthogonal pathways, Cytomarker enables prioritization of redundant markers for removal. Finally, if the user would like to find alternatives for a given marker in scenarios such as antibody unavailability, Cytomarker can quantify the ability of different markers to act as surrogates for efficient substitution.

To translate the marker list into an actionable panel, Cytomarker includes flexible functionality to identify validated reagents to create panels of interest. The built-in antibody explorer lets users browse antibodies matching their panel from a large and growing database of antibodies. This includes metadata such as validated applications to enable the user to filter their antibody selection based on a specific application or tissue type (e.g. flow vs. imaging, fixative type, organism of interest). In addition, users may download an html report that includes all figures, scores, and antibody information from the session that can be saved and shared for iterative panel refinement.

As a proof-of-concept experiment, we utilized Cytomarker to develop an antibody panel for highly multiplexed profiling of health mammary tissue. Specifically we sought to identify canonical and novel markers delineating luminal (inner ductal epithelium), basal (outer ductal epithelium) and fibroblast (surrounding stroma) cell populations. This interactive approach identified established markers for basal (KRT5, KRT14) and luminal (KRT8, KRT18) populations, as well as to our knowledge novel subpopulation markers including CLDN3 in luminal, COL17A1 in basal, APOD in fibroblasts, and ColIV in endothelial cells (**Fig. 1B-C** and **S. Table 1**). Next, we developed the antibody panel for these targets and conjugated the antibodies to heavy metals to perform highly multiplexed imaging using Imaging Mass Cytometry (IMC) on a healthy human breast tissue section (**Methods**). Following standardized data processing, single-cell quantification, and clustering, we reconstructed the cell type specific expression from the IMC data (**Fig. 1D**). Importantly, this validated the single-cell protein expression profiles of the four novel subpopulation markers identified from the scRNA-seq, though APOD was also present in immune populations, which may be due to difficulties segmenting immune cells from multiplexed imaging data^18^. Finally, we examined the spatial distribution of markers on a region of tissue where luminal, basal, and fibroblast populations were clearly distinct (**Fig. 1E**). As expected, the Cytomarker-derived novel subpopulation markers showed the expected tissue localization for newly identified markers (**Fig. 1F-H**), validating the overall approach.

To more broadly validate our marker prediction and this approach, we developed a novel antibody screening strategy to simultaneously validate RNA-based predictions of > 200 cell surface markers (**Fig. 2A**). Arrayed peripheral blood mononuclear cells (PBMCs) were labelled with a unique cell surface barcode and an experimental surface marker of interest, pooled, and stained with 12 lineage markers and secondary antibodies that targeted the unique experimental antibody from each well.

**Figure 2.**
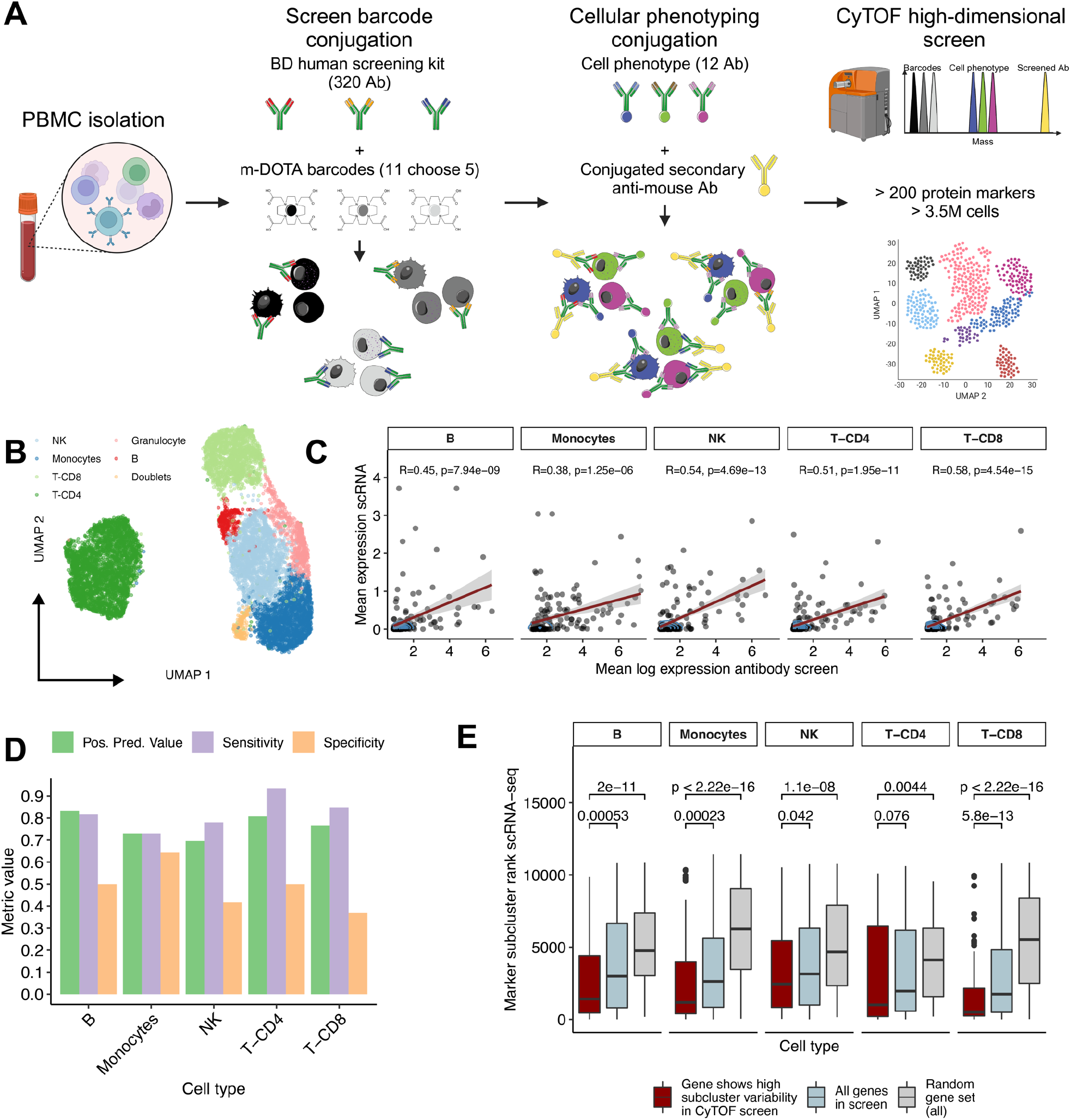
**A**. Schematic of high-dimensional antibody screen including (1) PBMC isolation, (2) conjugation of 300-plex screen barcodes, (3) conjugation of cellular phenotype antibodies, and (4) high-dimensional single-cell analysis with CyTOF. **B** UMAP of a cell subsample using 12 cell phenotype markers. **C** Cell-type-specific correlation between mean expression of antibody screen and scRNA-seq. **D** Positive predictive value, sensitivity, and specificity of using scRNA-seq to identify protein expression biomarkers from screen. **E** Highly-variable markers from antibody screen are ranked highly as scRNA-seq subcluster markers compared to control gene sets.

Analysis of cytometry by time-of-flight (CyTOF) measurement of >3.5m cells (**Methods**) deconvolved 6 broad immune populations (**Fig. 2B, S. Fig. 1**) and their labelling by all 220 surface markers including expected cell subset specific marker expression (**S. Fig. 2**).

Next, we broadly correlated cell type specific mRNA expression from scRNA-seq with the inferred protein expression from the antibody screen. We retrieved annotated PBMC scRNA-seq data^19^ and subset to cell types identified by the immune profiling panel in the CyTOF screen (**Methods**). We contrasted the mean expression of each of the 220 cell surface markers between the RNA and antibody screen, finding correlations between 0.38 and 0.58 and all p-values < 10^−5^ (**Fig. 2C**). Next, given marker selection can be framed as a prediction problem, we discretized each marker into high or low expression for each assay and assessed the ability of the scRNA to predict cell-type specific highly expressed markers in the CyTOF single cell antibody screen (**Methods**). Across cell lineages, this revealed high positive predictive value and sensitivity with moderate specificity (**Fig. 2D**), consistent with scRNA-seq being better powered to detect highly expressed genes. Incorporating orthogonal data of paired proteomic and RNA data from similar cell types^20^ and removing genes with low RNA-protein correlation markedly improved overall specificity (**S. Fig. 3** and **Methods**).

Finally, we investigated whether this platform could validate scRNA-seq predictions of granular sub- population markers. We reasoned that proteins exhibiting highly variable expression within cell lineages from the CyTOF screen were likely to be marker genes for cell subclusters, we ranked genes by their ability to discriminate scRNA-seq populations and contrasted these with the highly variable proteins from the screen. Across all lineages, highly variable markers from the screen were more likely to rank highly as scRNA-seq subcluster markers than other genes both in the screen and compared to a random gene set of the same size (**Fig. 2E** and **Methods**). Together, these results demonstrate the ability of our antibody screening approach to validate granular subpopulation markers found using scRNA-seq and the accuracy of Cytomarker designed antibody panels.

Despite our validation of Cytomarker, there are several limitations that can serve as a foundation for future development. The utility of an RNA based assay to derive antibody panels is limited by RNA-protein correlations that vary across genes and tissues^13^. Future versions may incorporate explicit information on these correlations that can be derived from paired RNA and proteomic assays^20^.

Furthermore, scRNA-seq typically profiles only highly expressed transcripts, meaning resulting panels may miss lowly expressed but important targets such as transcription factors or functional markers measurable by proteomics. Finally, well known canonical markers that are validated to identify cell types may be missed by scRNA-seq directed design. Therefore, future versions of Cytomarker may automatically identify populations and suggest known markers even if poorly discriminative from the RNA.

## Methods

### Cytomarker inputs and preprocessing

Cytomarker is an R/Shiny application designed to facilitate interactive cytometry panel design using scRNA-seq data using a comprehensive GUI and flexible user-selected inputs. Users may upload single-cell data in SingleCellExperiment^16^, Seurat^15^, and AnnData^14^ formats. Cytomarker requires single-cell data in this format to contain a metadata column that stratifies cells based on a property such as a cell type or state annotation, experimental condition or disease subtype. For brevity below, we subsequently refer to any grouping of cells as a *cell type*, though within Cytomarker it may represent any partitioning of heterogeneity. Alternatively, users may select one of 24 pre-annotated datasets obtained from the Tabula Sapiens human cell atlas^21^ covering a wide range of tissues and cell types.

To balance application speed and marker accuracy, each dataset is subsampled to between 2-3k cells in a stratified manner to balance cell type presence which has previously been shown to impact downstream results^22^. Specifically, (i) cells belonging to any cell type with fewer than 20 cells were removed, (ii) cells belonging to a cell type with 20-100 cells were included as-is, and (iii) cells belonging to a cell type with more than 100 cells were subsampled so only 3000 / (# cell types) remained. In addition, the data was filtered to remove genes with fewer than 10 total counts across all cells profiled. Finally, Cytomarker removes any cell type with fewer than 2 counts total or representing less than 0.5% of the total number of cells.

### Gene and protein symbol handling

Cytomarker provides implicit gene conversion and detection functions that attempt to recognize multiple gene formats. By default, Cytomarker sets will display gene symbols; if users upload datasets with features of a different nomenclature (such as Entrez or Ensembl IDs), these IDs will be automatically converted to gene symbols before analysis using the grch38 human reference genome annotations from the org.Hs.eg.db package. Cytomarker also automatically parses any user-uploaded custom panel lists to provide lists of gene aliases, where Cytomarker can identify gene identifiers that may have alternative names that appear in the dataset. Gene aliasing may occur if users upload data with Entrez IDs and have these features converted to gene symbols prior to analysis. Cytomarker reports these matches to the user in tabular format.

### Initial marker panel creation

Cytomarker has two options for initial panel creation: a cell type aware procedure (default) or running geneBasis^6^ with default settings. For the former, Cytomarker performs differential gene expression analysis using scran’s findMarkers function^23^. Cytomarker then produces a list of initially recommended markers by:

1. Each cell type specific list of differentially expressed genes is filtered to retain only those over- expressed (i.e. logFC > 0) in a cell type
2. For each cell type, initially ceiling((requested panel size + 10) / n), where n is the total number of cell types being analyzed, genes are retained from the cell type specific over- expression lists from findMarkers, by ranking based on logFC
3. The cell types with the most remaining markers are selected (initially this is all of them)
4. The mean log fold change per marker is computed across all remaining cell types (since each gene may be a marker for more than one cell type)
5. The marker with the lowest logFC is then removed
6. Steps 3-5 are repeated until the desired panel size is achieved
7. It is possible that the above procedure returns “multimarkers” in the final panel, which are genes that are highly over-expressed in more than one cell type. In these cases, Cytomarker does not explicitly remove these multimarkers, as certain tissue types such as PBMCs may benefit from markers that are highly expressed in multiple cell types
8. At the end of the recursive trimming procedure, if a particular cell type has only genes that are considered multimarkers, Cytomarker will replace the multimarker with the gene with the highest logFC specific to that cell type

### Panel scoring

Cytomarker provides validation scores for each panel created using a multinomial logistic regression model with k = 10 folds using the nnet package. The model uses the list of genes in the generated panel to predict the cell type, computing the positive predictive value (PPV) scores per class by comparing the predicted label to the label provided by the user. Balanced accuracy across all classification classes is also computed using the ‘bal_accuracy_vec’ function from yardstick R package.

### Interactive marker selection

To suggest additional markers for a given cell type, Cytomarker retrieves the cached list of top differentially expressed genes between the current cell type vs. all others. By default, the top 50 genes for the cell type requested that are not in the current panel are suggested, ranked using the computeMinRank function from scran using the top putative targets for each cell type. Similar genes for marker substitution are ranked based on Pearson correlation to the selected gene. Redundant genes are detected by computing the Pearson correlation of expression, then scoring the genes with the findCorrelation function from the caret R package^18^ and removing the gene with the higher mean correlation to the other genes in the panel.

### Antibody database linkage

The open source version of Cytomarker hosted at https://www.github.com/camlab-bioml/cytomarker uses antibody data provided by the antibody registry (https://www.antibodyregistry.org/ accessed January 2022) with targets linked based on gene symbol. The hosted version of Cytomarker at https://www.cytomarker.ai incorporates proprietary information provided directly by antibody providers that incorporates further metadata including validated applications.

### IMC preparation and staining

Each selected antibody was conjugated to a lanthanide-series metal isotope of a unique mass for detection via IMC. Conjugations were performed using the MaxPar antibody labeling kit (Standard Biotools) and according to the MaxPar labeling protocol. A formalin-fixed, paraffin-embedded, disease-free human breast tissue sample was baked at 60°C for 60 minutes to improve tissue preservation. The sample was deparaffinized in 3 solutions of fresh xylene for 10 minutes each and rehydrated in a graded series of alcohol (ethanol: water 100:0 twice, 96:4, 90:10, 80:20, 70:30, 5 min each). To perform antigen retrieval, a Decloaking ChamberTM NxGen (Biocare Medical) was used to incubate sample in HIER buffer (10mM Tris Base, 1mM EDTA, pH 9.2) for 30 minutes at 90°C, and then allowed to cool at room temperature. Once cooled, blocking was performed by incubating sample at room temperature in a 3% BSA, 5% horse serum in TBS blocking solution for 60 minutes. Next, sample was incubated at 4°C, overnight, in a solution of primary, unconjugated antibody (Collagen) diluted in blocking solution, followed by 3×5 min washed in TBS and secondary antibody (anti-rabbit) incubation at room temperature for 60 minutes. After another 3×5 min round of washed in TBS in order to remove any unbound secondary antibody, sample was incubated at 4°C overnight in a cocktail of conjugated primary antibodies diluted in blocking solution. Finally, sample was stained using iridium (a DNA intercalator, at 1:1000 for 5 min) and washed 3×5 min in YBS before being dried onto the glass slide for IMC acquisition. All antibodies were used at a concentration of 1:100.

### IMC Acquisition

A panorama of the entire tissue section was generated using a built-in brightfield microscope on an XTi imaging system (Standard Biotools). According to the presence of epithelium (ducts/alveoli in breast), regions of interest were selected to be laser-ablated in a rastered pattern at 800Hz. The raw data was processed using the commercial acquisition software (Standard Biotools).

### IMC data analysis

Multiplexed data in mcd format was processed using integrated flexible analysis pipeline (ImcPQ) available at https://github.com/JacksonGroupLTRI/ImcPQ, which first involves conversion of files to TIFF format, followed by cell segmentation using Mesmer^24^ to create masks of single cell entities. Masks were then overlaid with individual channels of each antibody target, and the mean expression levels of markers and spatial features of single cells were extracted. An output file was produced containing the single cell expression of every panel marker summarized by mean pixel intensity. To process this raw output, a cutoff of the 99th percentile was implemented to remove outliers resulting from pixel hotspots during acquisition. The pixel count was then normalized by dividing each channel by its maximum pixel intensity value. The Scanpy package^25^ was used for heatmap and UMAP generation. Clustering was performed using Leiden as implemented in Scanpy using default parameters.

### Cell lines and PBMC isolation

HEK 293T cells were obtained from ATCC. Cells were maintained in DMEM supplemented with 10% heat inactivated fetal bovine serum and 1% Penicillin/Streptomycin (Wisent). PBMCs were extracted from buffy coats obtained from healthy human donors supplied by the Canadian Blood Services Blood4Research Program (CBSREB# 2018.020) using the Ficoll gradient centrifugation method as previously described^26^. All buffy coats used in this study were approved for use by the University of Toronto Research Ethics Board (Protocol no. 25841). After isolation PBMC were frozen in a 1:1 ratio of 5% Fetal Bovine Serum (FBS) in PBS and 20% dimethyl sulfoxide (DMSO) in FBS and maintained in liquid nitrogen until further use. The day before usage, PBMC were thawed by warming up vials to 37°C and then washing cells to remove any leftover DMSO before to be cultivated in RPMI media supplemented by 10% heat inactivated FBS and 1% Penicillin/Streptomycin (Wisent). All cells were maintained in humid atmosphere at 37°C and 5% CO2.

### mDOTA preparation

Isotopically pure or natural abundance single isotope rare earth metals were purchased from Trace Sciences International as previously described^27^. Briefly, Metals were resuspended in L-buffer (20mM sodium acetate, Thermo) to a final concentration of 20mM. Two molar equivalents of maleimido-mono-amide-DOTA (Cedarlane) were added to each 20mM metal solution. Obtained solutions were then further diluted in DMSO to obtain 100μM working stock stored at -20°C. To determine optimal barcoding concentration, a titration was performed using HEK 293T cells. Succinctly, cells were labeled with increasing concentration of m-DOTA (5nM, 10nM, 25nM, 50nM, 100nM and 250nM) and then run through CyTOF as described below to determine the minimal concentration barcoding 100% of the cells.

### Barcode masterplate generation and checkboard verification

Eleven lanthanide-conjugated mDOTA barcodes were used at the following concentration in the creation of a barcode master plate: Y-89 (12.5 μM), In115 (12.5 μM), La139 (12.5 μM), Ce140 (5 μM), Pr141 (12.5 μM), Nd146 (5 μM), Eu151 (2.5 μM), Tb159 (2.5 μM), Ho165 (2.5 μM), Tm169 (1.25 μM), and Lu179 (2.5 μM). Five out of nine distinct mDOTA barcode probes were pipetted by OT2 pipetting robot (Opentrons) into each well of the 384-barcode master plate with 7 μL for each probe. To test the accuracy of barcoding using this master plate combination, checkerboard pattern and column pattern tests were performed by adding cells with or without IdU (5-Iodo-2’- deoxyuridine, Fisher Scientific) staining in alternating patterns. Cells were previously fixed with 4% PFA for 15 min, and permeabilized for 30 min with 0.2% saponin. 45 μL of cell suspension were loaded into individual wells in a 384-well plate, and then 5 μL of barcodes were added from the master plate and incubated for 30 minutes. Three washes with 3% BSA in PBS were performed with OT2 protocols before pooling. After pooling, samples were stained for DNA using 125 nM of Cell-ID Intercalator Ir (Standard BioTools, Cat# 201192B) overnight at 4 °C. The next day, cells were washed twice with 1% BSA/PBS and then washed twice with Cell Acquisition Solution Plus (CAS+) (Standard BioTools, Cat# 201244). After the last wash, cells were counted using a hemocytometer. Samples were dispatched into tubes that eventually containing 8.0 × 10^5^ cells each. Samples were split in multiples tubes to prevent clogging and enhance resuspension efficiency during acquisition. Each tube was spun and then pellets were resuspended in 50 μL of CAS+. All tubes were then loaded on the carousel of the CyTOF XT (Standard BioTools) and resuspended at 1.2 × 10^6^ cells per tube.

### Antibody screening

For antibody screening, the screened antibodies have been purchased in the form of lyophilized antibody already dispatched in multiwell plate (BD Human CD Screening Plate Cat# 560747). PBMC were fixed with 4%PFA for 10min at RT and then permeabilized using saponin 0,2% for 20min. After washing 500K PBMC were dispatched into three 96 well plates. Lyophilized and arrayed plates of antibodies were reconstituted as per manufacturer recommendation. Using the OT2 pipetting robot, 20ul of antibody was transferred from the Ab screening plate to the plates containing the cells. Ab were incubated for 15min then a resuspension was performed using the OT2 and Ab were incubated for 15 more min. After a total of 30min of incubation, 100ul of PBS+BSA was added per well. Plates were then centrifuges and one more wash with PBS/BSA was performed. Following the PBS/BSA wash, a PBS wash was performed to remove BSA before barcoding. After the PBS wash, barcoding was performed. To this end 10ul of mix barcodes were transferred from the barcoding masterplate to the plates containing cells. Barcodes were incubated for 30min with an agitation after 15min. After barcoding, three PBS/BSA washes were performed. After the last wash, samples from all wells were pooled in a single tube.

### Staining

Cells were then stained with anti-mouse and anti-rat secondary antibodies conjugated to Nd148 and Pt194 respectively. Using anti-rat and anti-mouse allows to reveal all the antibodies from the screening plate using only two antibodies. After staining with secondary antibodies, two washes with PBS/BSA were performed. After washing, cells were stained, with a second panel of antibodies allowing for identification of the different cell populations from PBMC. This panel is composed of custom conjugated antibodies (**S. Table 2**) and antibodies from the Immune Profiling Kit 7 markers (Standard BioTools, Cat#201302). Custom antibodies were prepared using the Maxpar X8 Multimetal Labeling Kit (Standard BioTools) according to manufacturer’s instructions. After staining with the second panel, cells were washed twice using cell wash solution. Cells were then resuspended in wash solution containing 125 nM of Cell-ID Intercalator Ir (Standard BioTools Cat#201192B) overnight at 4°C. On the day of acquisition, cells were washed twice with PBS/BSA 1% followed by two washes with CAS+ solution. Cells were then filtered using a 35μm cell strainer and then counted using a hemocytometer. 1,2M cells were then aliquoted in 100μl of CAS+ solution in FACS tube prior to be run on CyTOF.

### CyTOF acquisition

12 tubes were loaded on the carousel of the CyTOF XT (Standard BioTools) at a time. The CyTOF XT was programmed to resuspend the 1,2 × 10^6^ cells per tube in 1 mL of CAS+ plus EQ6 calibration beads (Standard BioTools) and acquired all tubes in one batch with a medium cleaning cycle run between each tube. Multiple runs were performed until all cells have been acquired. The event rate during acquisition was maintained between 200 and 300 events/second.

### PBMC screen data analysis

After acquisition, all files generated by each acquisition were concatenated using Premessa R package. After concatenation, the concatenated file was debarcoded using Premessa R package and the same barcoding key as the one used to generate the master plate scheme. Debarcoding parameter chose were separation threshold equal to 0,2 and maximal mahaloinis distance to 20. To interpret the cell types from the antibody screen, the CyTOF data was first subset from the 3,530,697 debarcoded cells to 10,000 cells for computational feasibility. Dimensionality reduction was performed with principal component analysis (PCA) and clustering was performed using Seurat clustering with resolution 0.5, both as implemented using SingleCellTK^28^ with all other parameters default. Clusters were interpreted into cell populations on the basis of examination of the 12 common lineage markers (**S. Fig. 1**). To subsequently expand the annotation to all >3.5m cells, a multinomial regression model was trained (using the multinom function in R, all options default) to predict cell cluster from the 10k cell subset, with predictions subsequently made across the full set of cells. All subsequent analysis was performed on log(x+1) expression values.

### Linking scRNA-seq to screen

Annotated scRNA-seq of PBMCs^19^ were retrieved from CellXGene^29^ (June 19th 2024). Data was subset to a single donor (“P5”) to match the antibody screen and minimize batch effects and subsampled to 10,000 cells for computational feasibility. Cells were subset so existing annotations matched those in the screen, specifically containing the terms “natural killer”, “monocyte”, “CD8”, “CD4”, “regulatory”, “B cell”. All subsequent expression measurements were taken from the “X” assay, corresponding to log normalized counts.

Next, RNA and protein identifiers were manually matched (**S. Table 3**). CyTOF expression values were subject to a log(x+1) transformation prior to comparison to RNA to approximately match the RNA transformation. Each marker discretized to a 1 or 0 based on being above (1) or below (0) median expression in their respective modality. Paired mass spectrometry ground truth data were retrieved from the supplementary information of a recent publication^15^ (sheet “RNA-protein distribution”). Similarly, we defined agreement between RNA and protein levels if a given gene/protein was above/below median expression in both modalities. Marker variance for the CyTOF expression values was computed following a mean-variance stabilizing transformation using LOESS regression in line with existing work^30^ to reduce the impact of mean expression on expression variance.

## Data and code availability

Raw and processed data for IMC and PBMC screen is available on Zenodo with DOI 10.5281/zenodo.13891857. Code to reproduce the analyses in this paper is available at https://github.com/camlab-bioml/2023-cytomarkerpaper-analysis. Cytomarker can be accessed publicly through a shinyapps.io server at https://www.cytomarker.ai. Cytomarker source code is available at https://github.com/camlab-bioml/cytomarker.

## Contributions

Concept: HWJ, KRC. Software development: MW, MG, EP. Data analysis: MW, MG, KRC. Experiments: GA, SL. Research funding: HWJ, APM, KRC. Manuscript: MW, MG, EP, GA, SL, HWJ, KRC.

## Acknowledgements and funding

This work is funded by NSERC Discovery Grants RGPIN-2020-04083 to KRC and RGPIN-2021-03404 to HWJ, a CIHR project grant PJT-175270 to KRC, an NSERC Alliance grant ALLRP 570709-2021 to HWJ and KRC, and a Canadian First Excellence Research Fund (CFREF) Medicine by Design Grand Questions grant (MbDGQ-2021-03) to HWJ and APM. Both KRC and HWJ acknowledge support from the Canada Research Chairs program and the Canadian Foundation for Innovation. MG acknowledges support from the University of Toronto Data Sciences Institute. We are grateful to Canadian Blood Services’ blood donors, who made this research possible. Figure 2A made using Biorender.

## Ethics approval

The data generation and analysis in this manuscript is covered under REB 20-0155-E and 20-0178-E at Mount Sinai Hospital, Toronto ON, Canada, and University of Toronto Research Ethics Board (Protocol no. 35956) CBSREB 2023-019.

