## Supplementary Figures for "Interactive design and validation of antibody panels using single-cell RNA-seq atlases"

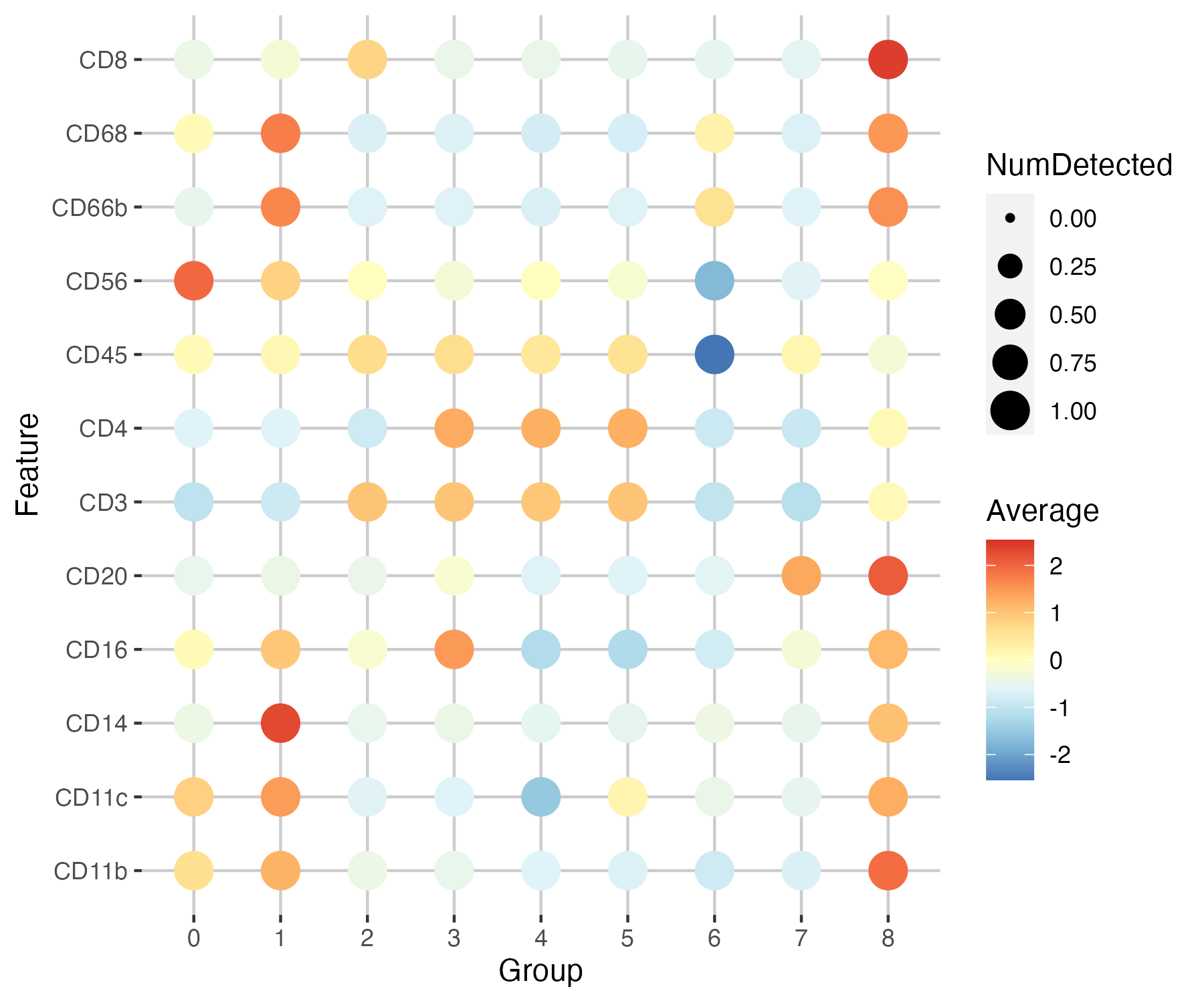

Supplementary Figure 1: Expression of the 12 common lineage markers by cluster for CyTOF screen.

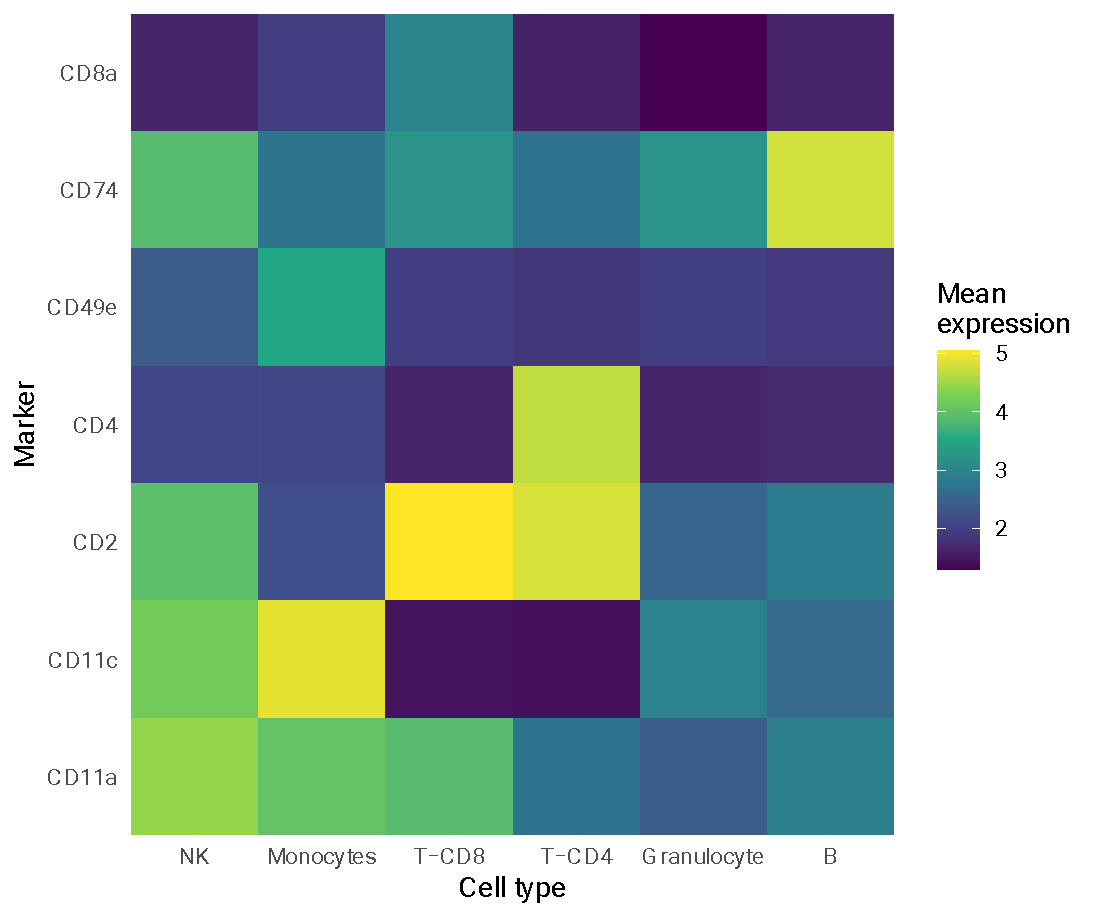

Supplementary Figure 2: Cell type specific mean expression of antibody library targets for a set of lineage markers.

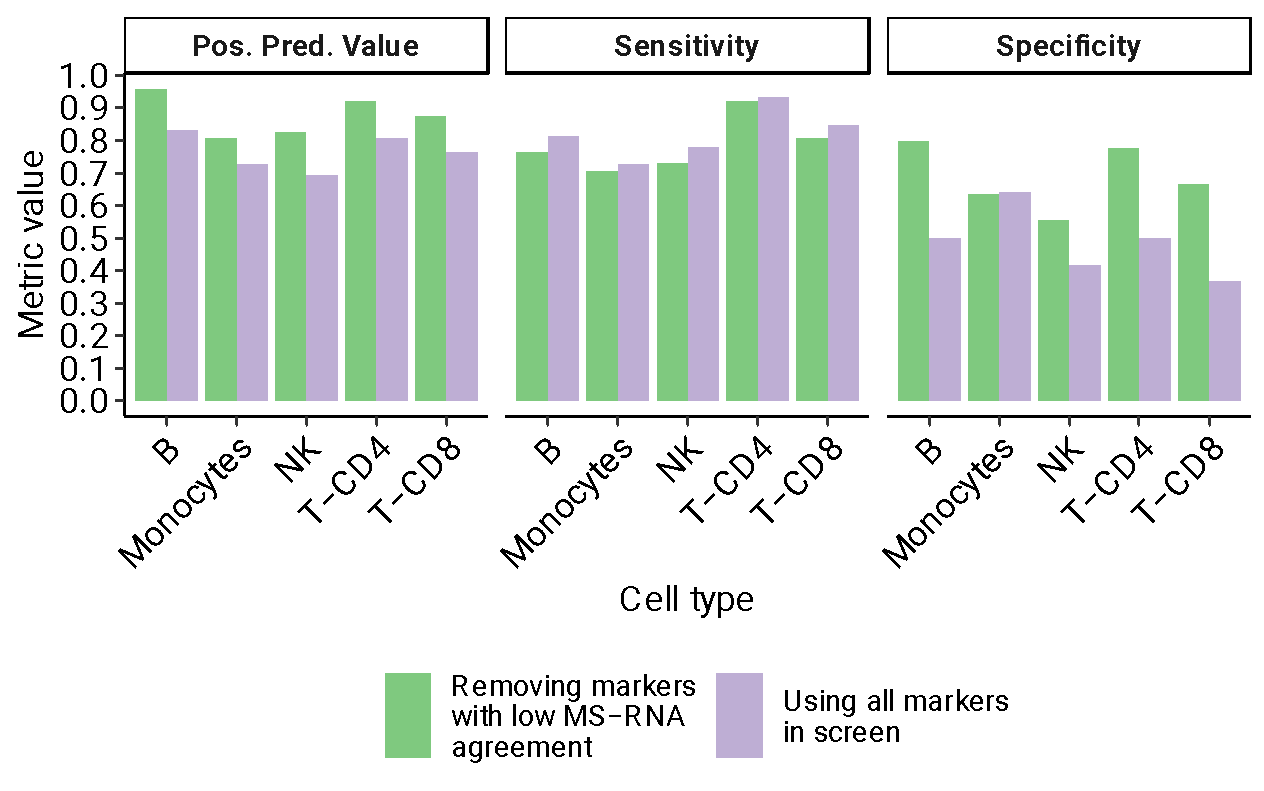

Supplementary Figure 3: Classification metrics across different major cell lineages with and without removing genes that show high expression correlation between protein as measured with mass spectrometry (MS) and RNA.
